# Assessing vector competence of mosquitoes from northeastern France to West Nile virus and Usutu virus

**DOI:** 10.1101/2023.02.07.527438

**Authors:** Jean-Philippe Martinet, Chloé Bohers, Marie Vazeille, Hubert Ferté, Laurence Mousson, Bruno Mathieu, Jérôme Depaquit, Anna-Bella Failloux

## Abstract

West Nile virus (WNV) and Usutu virus (USUV) are two arthropod-borne viruses that circulate in mainland France. Assessing vector competence has only been conducted so far with mosquitoes from southern France while an increasingly active circulation of WNV and USUV has been reported in the last years. The main vectors are mosquitoes of the *Culex* genus and the common mosquito *Culex pipiens*. Here, we measure the vector competence of five mosquito species (*Aedes rusticus, Aedes albopictus, Anopheles plumbeus, Culex pipiens* and *Culiseta longiareolata*) present in northeastern France. Field-collected populations were exposed to artificial infectious blood meal containing WNV or USUV and examined at different days post-infection. We show that (i) *Cx. pipiens* transmitted WNV and USUV, (ii) *Ae. rusticus* only WNV, and (iii) unexpectedly, *Ae. albopictus* transmitted both WNV and USUV. Less surprising, *An. plumbeus* was not competent for both viruses. Combined with data on distribution and population dynamics, these assessments of vector competence will help in developing a risk map and implementing appropriate prevention and control measures.

**Author summary:** West Nile virus (WNV) and Usutu virus (USUV) are on the rise in Europe and in France. WNV is reported in France as early as the 1960s in the Camargue and USUV more recently, in 2015 in eastern France. The re-emergence of WNV infections in the Camargue is associated with an expansion towards the North which is also favorable to maintain a viral transmission cycle. USUV frequently co-circulates with WNV sharing the same mosquito vectors. *Culex pipiens*, able to feed on birds and humans, is considered to be the main vector in France. Our study is the first to investigate the vector competence to WNV and USUV of five different mosquito species collected in northeastern France. We ascertain that French *Cx. pipiens* mosquitoes are competent to both WNV and USUV. More surprisingly, the mosquito *Aedes albopictus* from northeastern France was able to transmit WNV and USUV. Based on our result, we propose that surveillance of mosquitoes combined with viral detections must be implemented in northeastern France to allow early viral detection and timely intervention to prevent outbreaks of these two neurological diseases.

## Introduction

Since 2007, Europe has been facing an increase in local transmission of arboviral diseases with dengue virus (DENV), chikungunya virus (CHIKV), and Zika virus (ZIKV) transmitted by the invasive species *Aedes albopictus* in a human-to-human transmission cycle (1). Usutu virus (USUV) and West Nile virus (WNV) both belonging to the Japanese encephalitis (JE) serocomplex circulate primarily in an avian-mosquito cycle and are transmitted by *Culex* mosquitoes (2). Mammals (human, horses) can be infected but develop a low viremia, insufficient to infect mosquitoes; they are considered as dead-end hosts (3). WNV is regarded as the most important causative agent of viral encephalitis worldwide (4). Among the multiple WNV genotypes described up to date, lineages 1 and 2 have been associated with outbreaks in humans (5). On the other hand, USUV described under six lineages (6) circulates mainly among bird populations (mainly Passeriformes and Strigiformes) and human cases are rare. Both viruses are transmitted by *Culex* mosquitoes (7–9) and use a wide range of bird species as amplifying hosts.

In mainland France, WNV first emerged in 1962 (10) and was first isolated in *Culex modestus* mosquitoes in 1964 (11). USUV emerged in France in 2015 due to two different lineages originating from Germany and Spain (8); it was first isolated from blackbirds (*Turdus merula*) and *Culex* spp. mosquitoes (e.g. *Culex neavei, Culex pipiens, Culex perexiguus*, and *Culex perfuscus*) (8,9). While WNV causes outbreaks with severe neurological symptoms, only two cases of neuroinvasive USUV infections have been reported in humans (12,13). While the distribution and population dynamics of *Culex* mosquitoes are well documented, data on vector competence of French mosquitoes for WNV and USUV are incomplete. We know that *Culex modestus* and *Cx. pipiens* from the Camargue are competent for WNV (14,15) and no information is available for USUV. Here, we performed a vector competence analysis of mosquitoes (*Aedes rusticus, Ae. albopictus, Anopheles plumbeus, Cx. pipiens*, and *Culiseta longiareolata*) collected in northeastern France (inside a region bounded by Paris, Reims and Strasbourg) using experimental infections with WNV and USUV.

## Materials and Methods

### Ethic Statements

Animals were housed in the Institut Pasteur animal facilities accredited by the French Ministry of Agriculture for performing experiments on live rodents. Work on animals was performed in compliance with French and European regulations on care and protection of laboratory animals (EC Directive 2010/63, French Law 2013-118, February 6th, 2013). All experiments were approved by the Ethics Committee #89 and registered under the reference APAFIS#6573-201606l412077987 v2.

### Mosquito collections

*Aedes albopictus* Strasbourg were sampled from June to October 2022 using 31 ovitraps; 5782 eggs were collected. *Aedes rusticus, An. plumbeus, Cx. pipiens* and *Cu. longiareolata* were collected as immature stages in 2019-2022 (Table 1). Geographical distribution of mosquito sampling sites is detailed in Figure 1. The map was generated with RStudio v1.4.1103 (in combination with ggplot2 and ggspatial packages) (16–18). Eggs were immersed in water for hatching. Larvae and pupae were placed in pans containing 1 liter of dechlorinated water and a yeast tablet renewed every 2 days and maintained at 25±1°C. Pupae were collected in bowls placed in cages where adults emerged. Adults were fed with a 10% sucrose solution and kept at 28±1°C with a 12L:12D cycle and 80% relative humidity.

**Figure 1.**
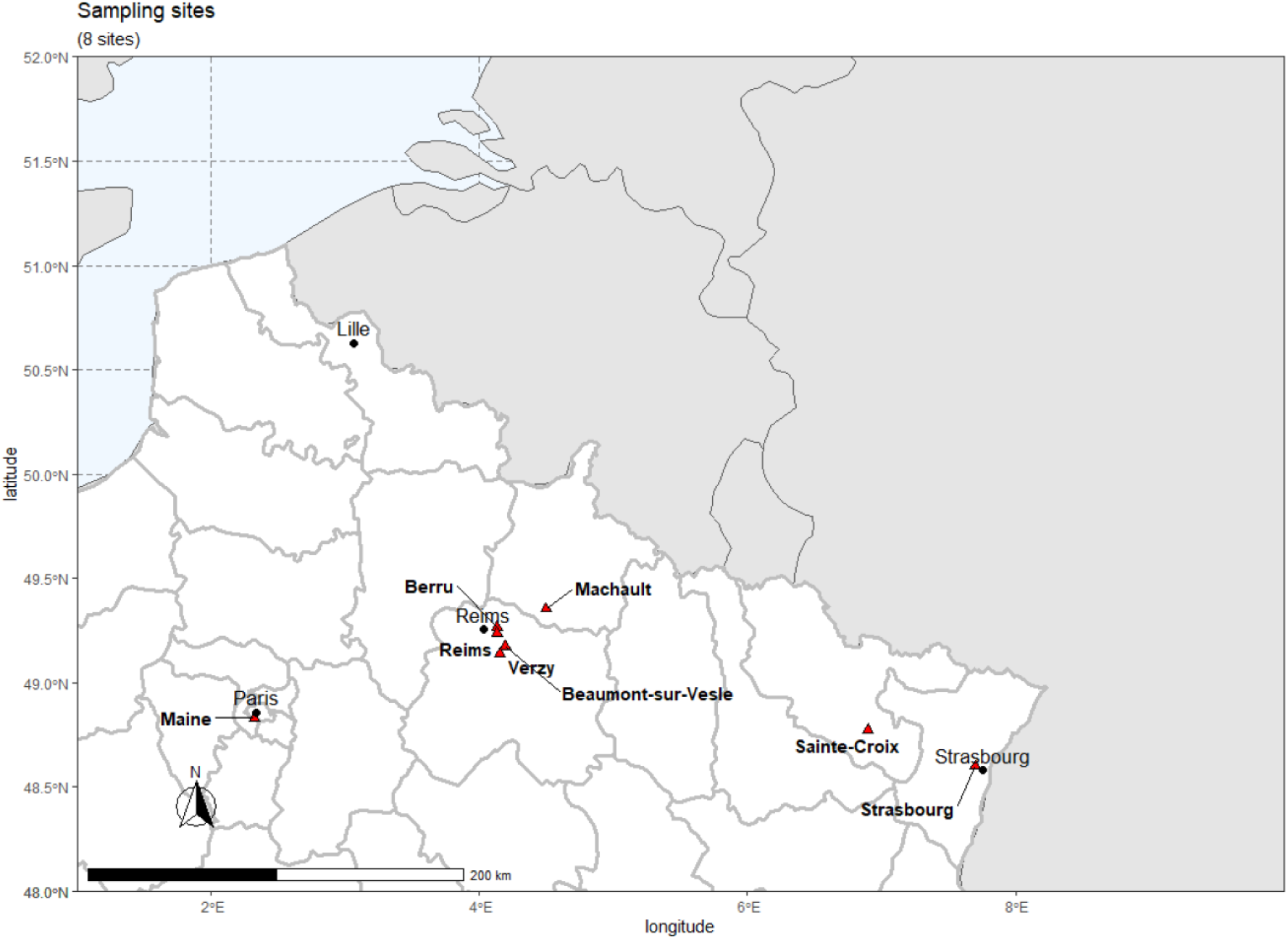
Map showing the different mosquito collections sites (red triangles) in northeastern France during the period 2019-2022.

**Table 1.**
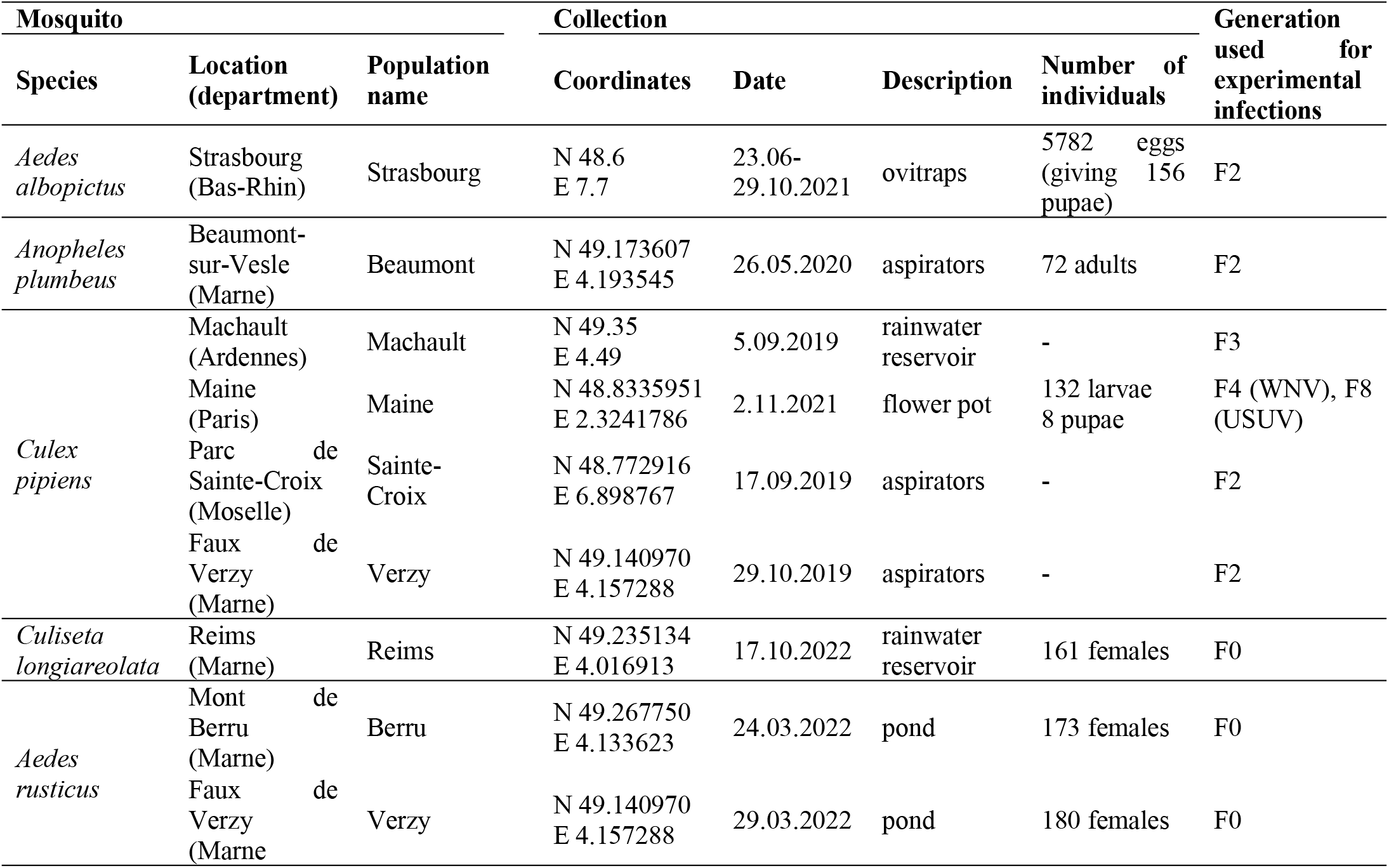
Mosquito species and populations collected in 2019-21 for assessing vector competence to WNV and USUV.

### Viral strains

The WNV-1a strain was isolated from a horse in Camargue (France) in 2000 (10). After 4 passages on Vero cells, the WNV stock was produced on *Ae. albopictus* C6/36 cells (19). The USUV Europa 3 strain (10214) was isolated in 2015 on Vero cells from the brain of a blackbird (8). All viral stocks were then produced on C6/36 cells and stored at −80°C until use.

### Mosquito infections and processing

Batches of 60 7-10-day-old females were transferred from cages into boxes and exposed to an infectious blood meal at a titer of 10^6.7^ plaque-forming unit (pfu)/mL for WNV and 10^7^ tissue culture infectious dose 50% (TCID_50_)/mL for USUV. The infectious blood meal containing 1.4 mL of washed rabbit erythrocytes, 700 μL of viral suspension and ATP at 1 mM as a phagostimulant, was put in a capsule covered with a pork intestine as membrane. This Hemotek feeding system was maintained at 37 °C. After 30 min of feeding, engorged mosquitoes were transferred in cardboard containers and supplied with 10% sucrose. Mosquitoes were maintained under controlled conditions (28±1°C, relative humidity of 80%, 12L:12D cycle) until examination. Mosquitoes were examined at different days post-infection (dpi) from 3 to 28 days depending on the number of fed mosquitoes and the mortalities observed after infection.

At a given dpi, surviving mosquitoes were cold anesthetized on ice. Then, legs and wings of each mosquito were removed and the proboscis was inserted into a pipette tip containing 5 μL of fetal bovine serum (FBS). After 30 min, the tip content was retrieved in 45 μL of L15 medium (Invitrogen, CA, USA). Then, head was isolated from abdomen and thorax. These two samples (head and thorax+abdomen corresponding to body) were separately ground in 300 μL of L15 supplemented with 2% FBS (Eurobio Scientific, Les Ulis, France), and centrifuged at 10,000×g for 5 min at +4 °C. Body (containing the midgut), head (possibly infected with viruses having disseminated from the midgut) were tested respectively for infection and dissemination while saliva was titrated to estimate transmission.

### Vector competence indices

To measure the vector competence, we used three indices which measured the role of the two main anatomical barriers in the progression of the virus in the mosquito after the infectious blood meal: (i) infection rate (IR) corresponding to the proportion of mosquitoes with infected midgut among mosquitoes exposed to the blood meal, (ii) dissemination rate (DR) referring to the proportion of mosquitoes having succeeded in disseminating the virus inside the mosquito general cavity among mosquitoes with infected midgut, and (iii) transmission rate (TR) which measures the proportion of mosquitoes with infectious saliva among mosquitoes having disseminated the virus. DR measures the efficiency of the midgut as a barrier to the dissemination of the virus inside the hemocele; the higher DR is, the less the midgut acts as a barrier to the dissemination of the virus. In addition to DR, TR measures the efficiency of the salivary glands as a barrier to the excretion of the virus in the saliva; as DR, the higher TR is, the less the salivary glands play the role of barrier to the transmission of the virus.

### Viral titration

Samples of saliva and extracts from bodies and heads of mosquitoes infected by WNV were titrated on Vero cells. Six-well plates containing confluent monolayers of Vero cells were inoculated with serial 10-fold dilutions of samples and incubated for 1 h at 37°C. Cells were then covered with an overlay consisting of DMEM (Gibco, CA, USA), 2% FBS, 1% antibiotic-antimycotic mix (Invitrogen, Gibco) and 1% agarose and incubated at 37°C. Cells were incubated 5 days. Lytic plaques were then counted after staining with a solution of safranine (0.5% in 10% formaldehyde and 20% ethanol). For mosquitoes infected by USUV, serial dilutions of saliva were inoculated on C6/36 cells in 96-well plates; each well was inoculated with 50 μL of diluted samples for one hour at 28 °C and after removing the inoculum, cells were covered with 150 μL of carboxymethylcellulose (CMC) supplemented with L-15 medium. After incubation at 28 °C for 5 days, cells were fixed with 3.6% formaldehyde, washed and hybridized with anti-flavivirus monoclonal antibody (catalog number: MAB10216, Millipore, CA, USA), and revealed by using a fluorescent-conjugated secondary antibody (catalog number: A-11029, Life Technologies, CA, USA), with dilution factors 1:200 and 1:1000, respectively. Foci were counted under a fluorescent microscope and titers were expressed as focus forming units (ffu)/sample.

### Statistical analysis

IR, DR and TR were compared using Fisher’s exact test and viral loads using Kruskal-Wallis test. Statistical analyses were conducted using the Stata software (StataCorp LP, Texas, USA). p-values < 0.05 were considered significant.

## Results

### Some populations of *Culex pipiens* are able to transmit WNV and USUV

To ascertain that *Cx. pipiens* mosquitoes were susceptible to WNV, we examined three populations for infection, dissemination and transmission at different dpi (Fig. 2A). We analyzed a total of 153 mosquitoes: 64 from Machault, 51 from Maine, and 38 from Verzy. When examining IR reflecting the success of midgut infection, we found that IRs were significantly higher at 10 dpi (77.8%, N=27 mosquitoes examined) than at 14 dpi (51.3%, N=37) for Machault (Fisher’s exact test: p=0.03). For Maine, IRs were significantly different at 3 dpi (44.4%, N=18), 7 dpi (66.7%, N=15), and 10 dpi (22.2%, N=18) (Fisher’s exact test: p=0.04). And, for Verzy, IRs were low: 0% (N=14) at 7 dpi and 8.3% (N=24) at 14 dpi (Fig. 2B). When measuring the viral load in bodies, mean values were similar at 10 dpi (10^6.7^ viral particles, N=21) and 14 dpi (10^6.6^, N=19) for Machault (Kruskal-Wallis test: p=0.22). For Maine, mean values significantly increased over dpi (Kruskal-Wallis test: p=0.008): 10^3.2^ (N=8) at 3 dpi, 10^4.4^ (N=10) at 7 dpi, and 10^5.2^ (N=4) at 10 dpi. For Verzy, the mean value was 10^5.3^ (N=2) at 14 dpi (Fig. 2E). When measuring DR translating the success of virus dissemination into the head, we found that DRs were similar at 10 dpi (33.3%, N=21) and 14 dpi (36.8%, N=19) for Machault (Fisher’s exact test: p=0.82). For Maine and Verzy, DRs were equal to 0% at all dpi examined (Fig. 2C). When assessing the viral load in heads, mean values were similar at 10 dpi (10^6.2,^ N=6) and 14 dpi (10^6.1^, N=7) for Machault (Kruskal-Wallis test: p=0.67). For Maine and Verzy, no viral particles were detected (Fig. 2F). When estimating TR describing the success of virus transmission, we detected that TRs were comparable between 10 dpi (14.3%, N=7) and 14 dpi (42.9%, N=7) for Machault (Fisher’s exact test: p=0.24). For Maine and Verzy, TRs were equal to 0% at all dpi examined (Fig. 2D). When estimating the viral load in saliva, mean values were similar at 10 dpi (10^5.3^, N=1) and 14 dpi (10^4.8^, N=3) for Machault. For Maine and Verzy, no viral particles were detected (Fig. 2G).

**Figure 2.**
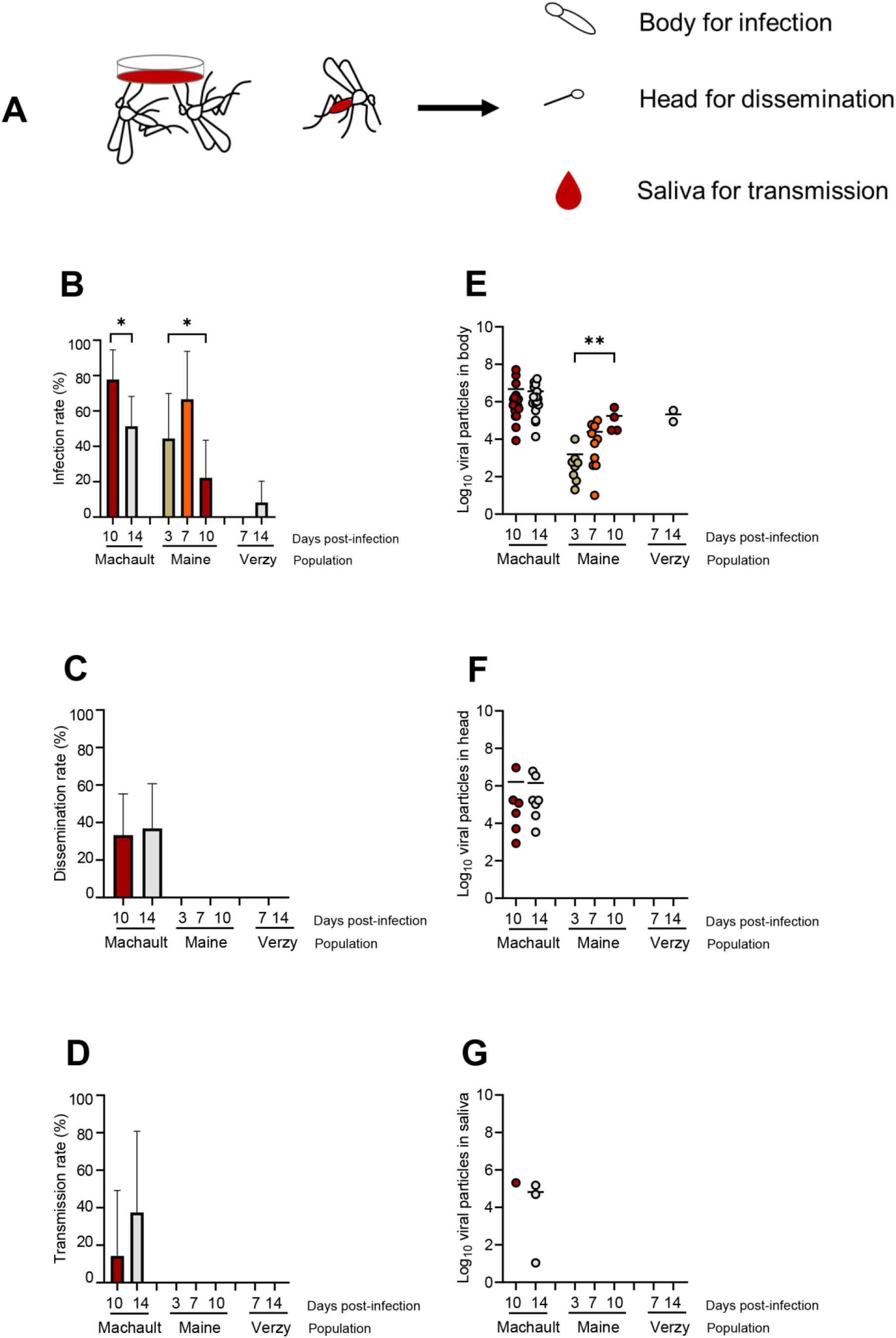
*Culex pipiens* (Machault, Maine and Verzy) experimentally infected with WNV provided at a titer of 10^6.7^ pfu/mL (A) and examined for infection (B), dissemination (C) and transmission, (D) at different days post-infection together with viral loads in body (E), head (F) and saliva (G). Infection rate refers to the proportion of mosquitoes with body infected among examined mosquitoes. Dissemination rate corresponds to mosquitoes with head infected (indicating a successful viral dissemination beyond the midgut barrier) among mosquitoes with infected body. Transmission rate indicates the proportion of mosquitoes presenting viral particles in saliva among mosquitoes with infected head. Stars indicate statistical significance of comparisons: *p≤0.05, **0.001≤p≤0.01.

To define whether *Cx. pipiens* was susceptible to USUV, we examined a total of 73 mosquitoes from Sainte-Croix at 7, 14, and 21 dpi. We found that IR was equal to 0% at 7 dpi (N=27) and 21 dpi (N=23) resulting in DR=0% and TR=0%. At 14 dpi, IR value was 4.3% (N=23) with one mosquito hosting 120000 viral particles in the body; this infected mosquito was able to disseminate (DR=100%) and to transmit (TR=100%) with 14 viral particles in the saliva (data not shown).

### *Anopheles plumbeus* is able to become infected but not to transmit WNV and USUV

To verify that as expected, *Anopheles* mosquitoes are not susceptible to arboviruses, we examined 46 *An. plumbeus* exposed to WNV and 60 to USUV.

For WNV, when examining IR, we found that IRs were comparable at 7 dpi (23.8%, N=21), 14 dpi (31.8%, N=22), and 21 dpi (23.5%, N=17) (Fisher’s exact test: p=0.79; Fig. 3A). The mean values of viral load in bodies were similar at 7 dpi (10^6.6^, N=5), 14 dpi (10^6.3^, N=7) and 21 dpi (10^6.4^, N=4) (Kruskal-Wallis test: p=0.79; Fig. 3D). When measuring DR, we found that DR value was equal to 0% at 7 dpi (N=5) and values were similar between 14 dpi (14.3%, N=7) and 21 dpi (50%, N=4) (Fisher’s exact test: p=0.40; Fig. 3B). The mean values of viral load in heads were similar between 14 dpi (10^5.0^, N=1) and 21 dpi (10^6.2^, N=2) (Fig. 3E). When estimating TR, we found that TRs were equal to 0% (Fig. 3C) and no viral particles were detected (Fig. 3F).

**Figure 3.**
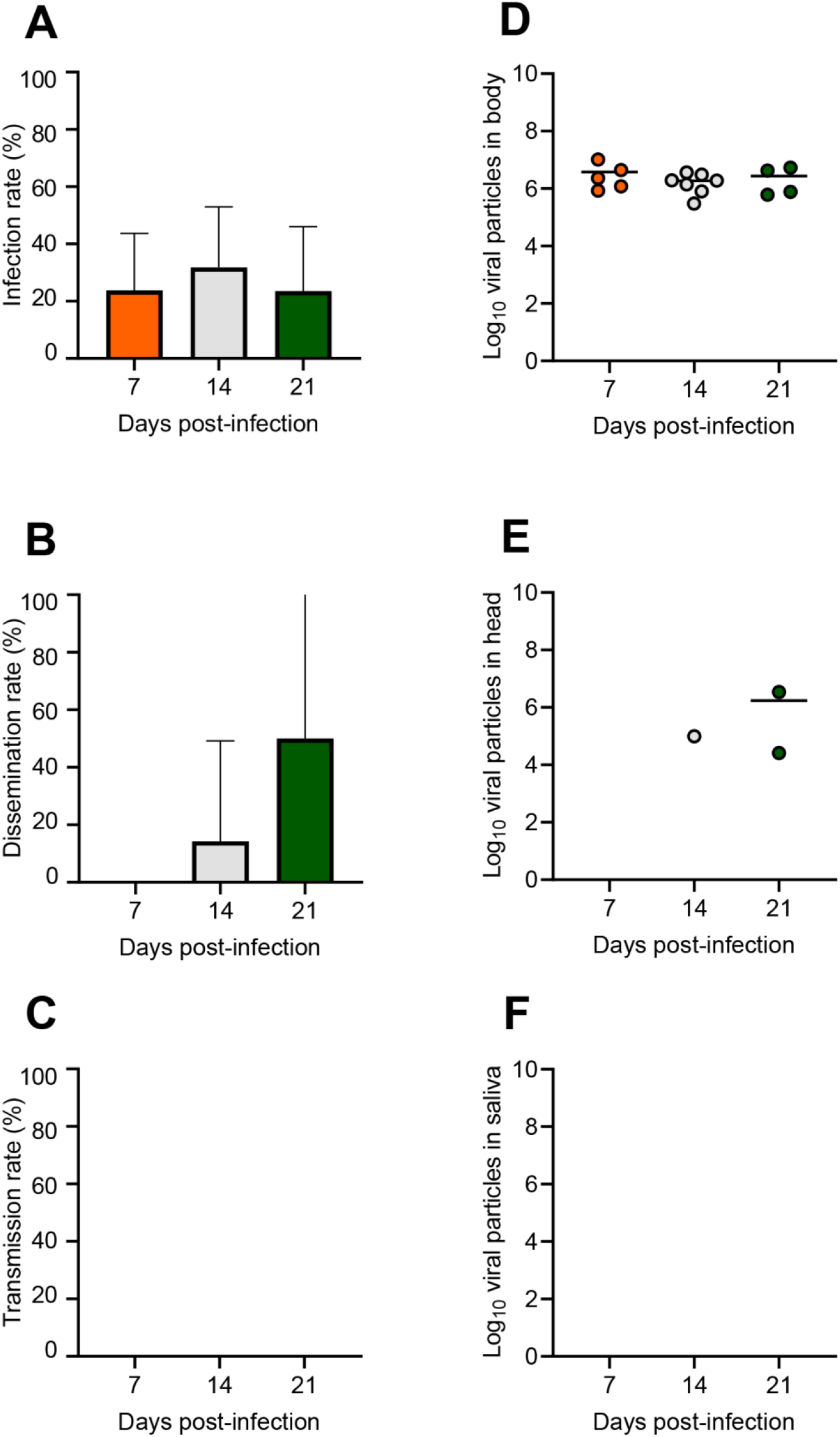
*Anopheles plumbeus* (Beaumont) experimentally infected with WNV provided at a titer of 10^6.7^ pfu/mL and examined for infection (A), dissemination (B) and transmission (C) at different days post-infection (7, 14, and 21) together with viral loads in body (D), head (E) and saliva (F).

For USUV, infection of bodies was only detected at 14 dpi (IR=4.35%, N=23) while IR was 0% at 7 dpi (N=23). No dissemination (DR=0%) or transmission (TR=0%) was detected (data not shown).

### *Aedes rusticus* is able to transmit WNV but not USUV

To search for secondary vectors of WNV and USUV beside the main vector *Cx. pipiens*, we examined 30 *Ae. rusticus* mosquitoes from Berru exposed to WNV and 51 mosquitoes from Berru and 45 from Verzy exposed to USUV. For WNV, we found that at 7 dpi, IR was 50% (N=30), DR of 33.33% (N=15), and TR of 40% (N=5) (Fig. 4A). The mean numbers of viral particles were 10^7.05^ viral particles in bodies (N=15), 10^6.6^ in heads (N=5), and 37 in saliva (N=2) (Fig. 4B). For USUV, infection (IR=0%), dissemination (DR=0%), and transmission (TR=0%) were not detected at 4, 7, and 10 dpi for Berru and Verzy populations (data not shown).

**Figure 4.**
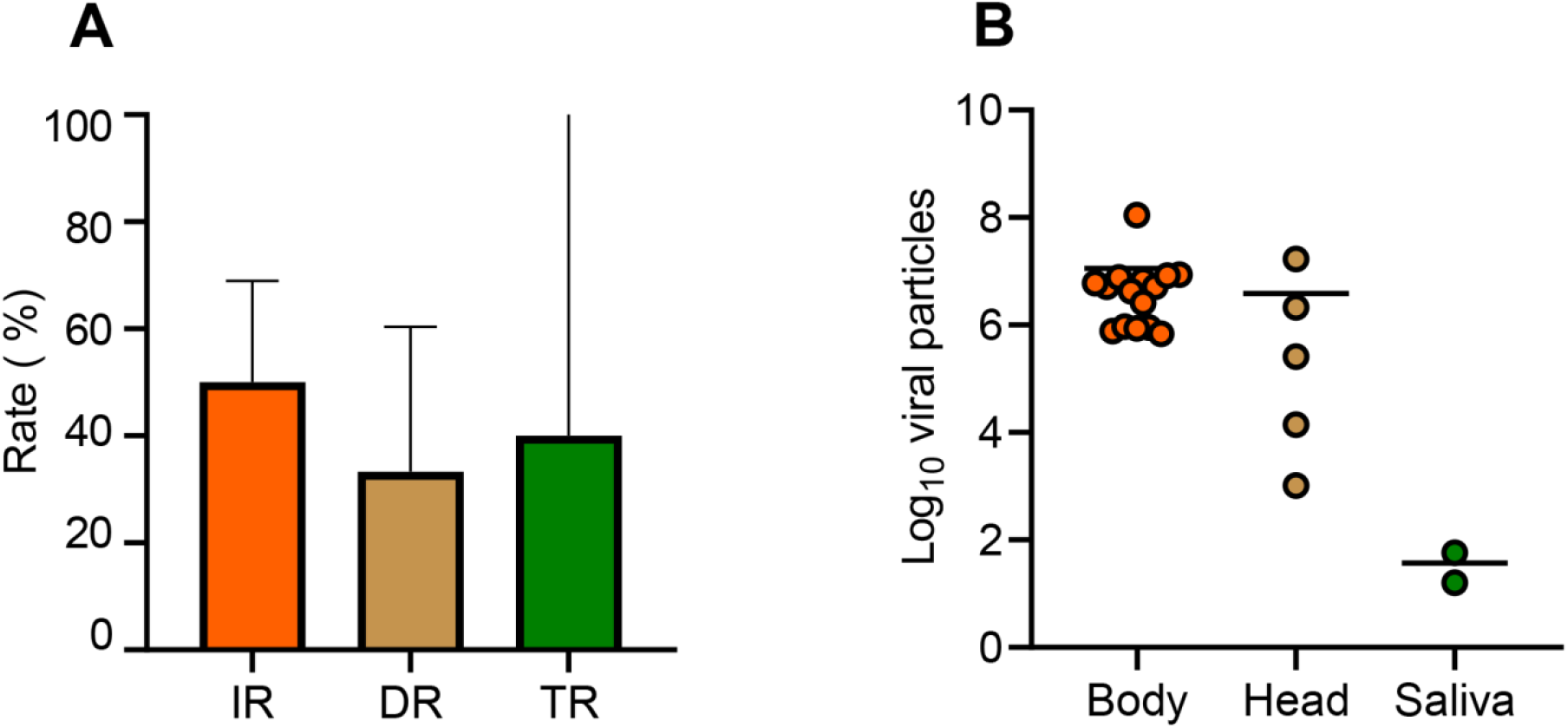
*Aedes rusticus* (Berru) experimentally infected with WNV provided at a titer of 10^6.7^ pfu/mL and examined at 7 days post-infection for infection, dissemination and transmission (A), and viral loads in body, head and saliva (B).

### *Aedes albopictus* transmits WNV and to a lesser extent, USUV

To see if this invasive mosquito could transmit *Culex*-borne viruses such as WNV and USUV, we examined 156 *Ae. albopictus* mosquitoes exposed to WNV and 150 to USUV.

For WNV, when examining IR, we found that IRs were similar at 3 dpi (20.8%, N=24), 7 dpi (25%, N=24), 10 dpi (29.2%, N=24), 14 dpi (29.2%, N=24), 17 dpi (54.2%, N=24), 21 dpi (41.7%, N=24), and 28 dpi (8.3%, N=12) (Fisher’s exact test: p=0.07; Fig. 5A). Mean values of viral load in bodies were comparable at 3 dpi (10^4.07^, N=5), 7 dpi (10^5.6^, N=6), 10 dpi (10^7.2^, N=7), 14 dpi (10^7.8^, N=7), 17 dpi (10^7.5^, N=13), 21 dpi (10^8.0^, N=10), and 28 dpi (10^7.3^, N=1) (Kruskal-Wallis test: p=0.22; Fig. 5D). When measuring DR, we found that DR was equal to 0% (N=5) at 3 dpi, increased to 33.33% (N=6) at 7 dpi, 100% (N=7) at 10 dpi, 100% (N=7) at 14 dpi, and dropped to 61.5% (N=13) at 17 dpi et 80% (N=10) at 21 dpi (Fisher’s exact test: p=0.001; Fig. 5B). Mean values of viral load in heads were similar at 7 dpi (10^4.5^, N=2), 10 dpi (10^6.5^, N=7), 14 dpi (10^7.3^, N=7), 17 dpi (10^5.9^, N=8), 21 dpi (10^5.3^, N=8), and 28 dpi (10^5.8^, N=1) (Kruskal-Wallis test: p=0.65; Fig. 5E). When estimating TR, we found that TRs were equal to 0% (N=2) at 7 dpi, 42.8% (N=7) at 10 dpi, 100% (N=7) at 14dpi, 62.5% (N=8) at 17 dpi, 75% (N=8) at 21 dpi, and 100% (N=1) at 28 dpi (Fisher’s exact test: p=0.072; Fig. 5C). Mean values of viral load in saliva were similar at 10 dpi (10^2.9^, N=3), 14 dpi (10^2.3^, N=7), 17 dpi (10^3.3^, N=5), 21 dpi (10^3.3^, N=6), and 28 dpi (10^1.5^, N=1) (Kruskal-Wallis test: p=0.52; Fig. 5F).

**Figure 5.**
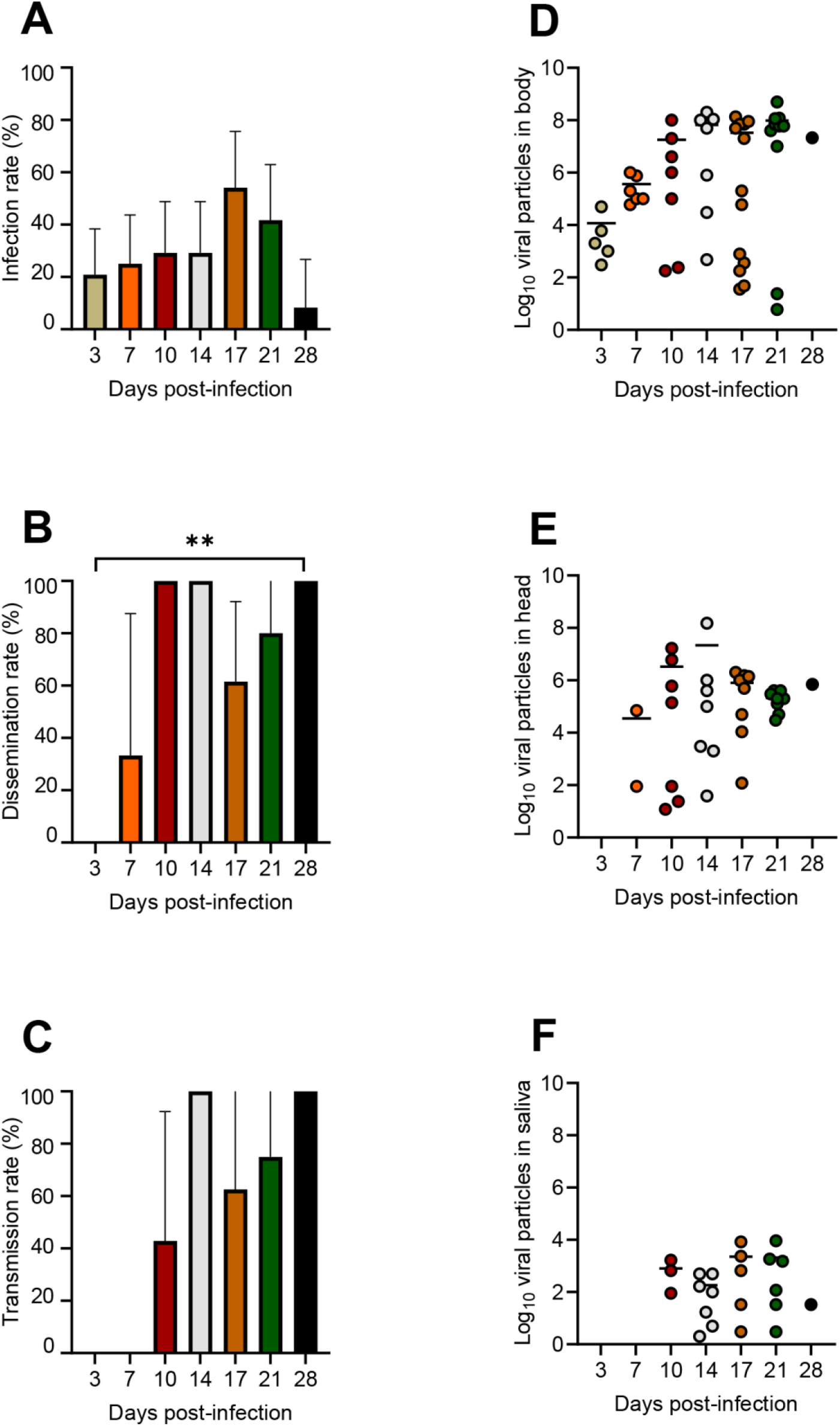
*Aedes albopictus* (Strasbourg) experimentally infected with WNV provided at a titer of 10^6.7^ pfu/mL and examined for infection (A), dissemination (B) and transmission (C) at different days post-infection (3, 7, 10, 14, 17, 21, and 28) together with viral loads in body (D), head (E) and saliva (F). Stars indicate statistical significance of comparisons: **0.001≤p≤0.01.

For USUV, we only examined the transmission. TRs were equal to 4.2% (N=24) at 10 dpi, 29.2% (N=24) at 14 dpi, 12.5% (N=24) at 17 dpi, 16.7% (N=24) at 21 dpi, and 16.7% (N=6) at 28 dpi (Fisher’s exact test: p=0.01; data not shown). Mean values of viral load in saliva were 10^0.8^ (N=1) at 10 dpi, 10^2.2^ (N=7) at 14 dpi, 10^2.4^ (N=3) at 17 dpi, 10^2.0^ (N=4) at 21 dpi, and 10^3.2^ (N=1) at 28 dpi (Kruskal-Wallis test: p=0.26; data not shown).

### *Culiseta longiareolata* is not able to transmit USUV

Due to the difficulty in feeding *Cs. longiareolata* mosquitoes in BSL-3 conditions and keeping them alive, only 8 individuals were examined at 10 dpi. None of them were able to transmit USUV (data not shown).

## Discussion

We show that some field-collected mosquitoes from northeastern France are competent vectors of WNV and USUV; WNV was transmitted by *Cx. pipiens*, *Ae. rusticus*, and *Ae. albopictus* and USUV was transmitted by *Cx. pipiens* and *Ae. albopictus*.

*Aedes albopictus* is an invasive mosquito species introduced in mainland France in 1999 (20) and is now established in 67 out of 96 departments (21). It is a known vector of multiple arboviruses such as DENV or CHIKV (22). French populations of *Ae. albopictus* are experimentally competent to DENV, CHIKV and ZIKV (23,24). We show that *Ae. albopictus* from Strasbourg was able to transmit WNV from 10 days post-infection. This length of the extrinsic incubation period (EIP) is close to the value estimated for Italian mosquitoes (i.e. 9-14 days; (25)). We also found that *Ae. albopictus* was able to transmit USUV from 10 days post-infection with 10^0.8^ viral particles detected in saliva of one mosquito. Combined with the detection of USUV in field-collected *Ae. albopictus* (26,27), our results on vector competence for USUV, are in favor of a role of *Ae. albopictus* in the transmission cycle of USUV (28,29).

*Aedes rusticus* is ubiquitous in northern France and present in 12 countries of western Europe (30). The species mainly present in forested environments in close contacts with avian populations is a human biting mosquito, active from April to August (Martinet, personal communication). We show that viral transmission occurs at 7 dpi and certainly before, indicating a high potential of *Ae. rusticus* to behave as competent WNV vector on the field. This species is the fourth referenced *Aedes* species of northeast Europe to be competent to WNV beside *Ae. caspius, Ae. detritus* and *Ae. japonicus* (14,31,32). However, no WNV-infected mosquitoes were detected on the field (33).

*Anopheles plumbeus* was one of the historical malaria vectors in northern Europe (34). This mosquito is present from late spring to the end of September, and females feed mostly on mammals (35). This species had been described as competent for the WNV lineage 2 (36). We found that *An. plumbeus* became infected but was not able to transmit WNV lineage 1a. More studies are required to clarify the vector status of *An. plumbeus* as this species is actively colonizing anthropic biotopes (37). Screenings performed on *An. plumbeus* in Germany in 2007-2008 showed no evidence of WNV-infected specimens (38).

*Culex pipiens* populations used in this study were collected in different biotopes. *Culex pipiens* Machault from the department of Ardennes was collected in a rural area, at proximity of human habitations and domestic animals. The Verzy population from the department of Marne was sampled in a sylvatic environment often visited by hikers. The Sainte-Croix population in the department of Lorraine was collected in a zoological park at a short distance from *Strix nebulosa* enclosure and Maine population from Paris was collected in an urban environment. We provide one of the first vector competence data on *Culex* populations from northern France in addition to data for mosquitoes from southern France (14,15). Populations from West Europe (Germany, The Netherlands, Switzerland, United-Kingdom) were experimentally capable to transmit WNV (39–42). Furthermore, circulation of WNV in Germany has been reported in 2018 (43). We found that Machault population was able to transmit WNV but not Maine and Verzy highlighting variations of vector competence depending on the mosquito population in addition to the virus genotype (44). We determined that Machault population was composed of the two biotypes, *Cx. pipiens pipiens* and *Cx. pipiens molestus*, and hybrids *pipiens*/*molestus* (S1 Table). The two forms have distinct host preferences (*Cx. pipiens pipiens* biting mainly birds and *Cx. pipiens molestus*, mainly mammals, especially humans) (45). Therefore, hybrids could ensure the transmission of the virus from the bird to the human.

One *Cx. pipiens* Sainte-Croix was able to transmit USUV at 14 days post-infection with 14 viral particles detected in the saliva. Our result is on the line with the vector competence described for local populations of *Cx. pipiens* in Germany and Switzerland (41,42). It is also correlated with the different episodes of USUV infections of *Strix nebulosa* and *Strix aluca* in zoological parks in the last years (46). Therefore, monitoring of mosquito populations in zoological parks housing susceptible bird species could be of help to prevent USUV circulation.

*Culiseta longiareolata* is an ornithophilic species widely distributed in the southern Paleartic region (47). These last years, it has been spreading to western Europe including several countries bordering France such as Belgium and Germany (47,48). This species is considered as a vector of blood parasites in birds and is not likely to feed on humans (35). Despite the limited number of mosquitoes examined, we show that *Cs. longiareolata* from Reims was not able to transmit USUV.

WNV-competent birds mainly belong to the Passeriformes and Charadriiformes (7). Introduction of WNV in mainland France is believed to be done by migrating birds and main migratory routes crosses France from southwest to northeast (49). The viraemia in birds does not last long enough (4-5days) to cover the duration of the migration (15-20 days from sub-Saharan Africa to Europe). Therefore, the probability that an WNV-infected bird arrives in Europe directly from sub-Saharan Africa is low (50) and secondary spots of viral contaminations are suspected in resting sites for migrating birds, which are numerous in northeastern France.

WNV and USUV share the same hosts in their transmission cycle: birds as amplifying hosts and *Culex* mosquitoes as vectors. An increasing circulation of USUV would have important implications on WNV as it affects significantly human health. USUV may have a strong cross-neutralizing potential towards WNV; the two closely related flaviviruses of the JEV serocomplex cannot persist in the same ecological niche due to cross-protective avian herd immunity (51). Furthermore, *Cx. pipiens* mosquitoes appear to be a major vector for both WNV and USUV in Europe. This species feeds on birds and humans which then, can act as a bridge vector for spillovers from birds to humans and horses (52). USUV and WNV outcompete in mosquitoes indicating that the chance of concurrent USUV and WNV transmission *via* a single mosquito bite is low (53).

Differences of climate could also be considered in the disparity of WNV circulation in the country between north and south of France. Warm Mediterranean climate in the south could enhance virus circulation by: (i) contributing to increase mosquito densities and intensify contacts with bird hosts, and (ii) shortening the EIP allowing WNV to be transmitted earlier in the south than in the north. In the context of global change, warmer summers and autumns can decrease the temperature differential between North and South of France, expanding the geographical area and the period of viral contamination.

In conclusion, we report two new putative vectors for WNV in northeastern France (*Ae. albopictus* and *Ae. rusticus*) aside from its known vector *Cx. pipiens*. WNV and USUV transmission overlaps sharing the same hosts and highlighting the importance of further studies on the interactions between the two viruses within vertebrate hosts and vector populations.

## Acknowledgments

The authors are grateful to Eva Nast for field collection of mosquitoes, and André Yébakima and Pierre Manuellan for collecting *Culex* mosquitoes in Paris 14^th^ district. We warmly thank Dr Jennifer Lahoreau for sampling at Parc de Sainte-Croix zoological park. We also thank Benjamin Dupuis for helping in genotyping *Culex pipiens* mosquitoes.

## Authors contribution

**Conceptualization:** Jean-Philippe Martinet, Anna-Bella Failloux

**Formal analysis:** Chloé Bohers, Marie Vazeille, Laurence Mousson,

**Funding acquisition:** Jérôme Depaquit, Anna-Bella Failloux

**Investigation:** Jean-Philippe Martinet, Chloé Bohers, Marie Vazeille, Laurence Mousson, Bruno Mathieu, Hubert Ferté, Jérôme Depaquit

**Writing – original draft:** Jean-Philippe Martinet, Anna-Bella Failloux

**Writing – review & editing:** Jean-Philippe Martinet, Marie Vazeille, Anna-Bella Failloux

---

**S1 Table.** Molecular assay based on indels in the flanking region of a microsatellite locus CQ11 to distinguish the two forms of *Culex pipiens, pipiens and molestus*. The method is described in Bahnck & Fonseca. 2006 (Rapid assay to identify the two genetic forms of Culex (Culex) pipiens L. (Diptera: Culicidae) and hybrid populations. Am J Trop Med Hyg. 2006 Aug;75(2):251-5). One mosquito leg was placed in a tube containing the PCR mix composed of two antisense primers at a final concentration of 0.15 μM, the sense primer at 0.25 μM, buffer (1X), dNTPs at 250 μM, MgCl2 at 1.5 mM, BSA (Bovine serum Albumin) at 0.135μg/μL, one unit of Taq polymerase and 5 μL of the DNA extract. The primers used were: pipCQ11R 5’-CATGTTGAGCTTCGGTGAA-3’, molCQ11R 5’-CCCTCCAGTAAGGTATCAAC-3’ and CQ11F2 5’-GATCCTAGCAAGCGAGAAC-3’. The amplification program started with 15 min at 94°C, 35 cycles of 94°C for 30s, 54°C for 30s and 72°C for 40s and finally a 5 min elongation phase at 72°C. PCR products were separated by electrophoresis on a 2% agarose gel. P, *Cx. pipiens pipiens;* M, *Cx. pipiens molestus;* PM, hybrids *pipiens/molestus*

| Population | Machault | Maine | Sainte-Croix | Verzy |
| --- | --- | --- | --- | --- |
| Number of mosquitoes tested | 19 | 16 | 16 | 17 |
| Pipiens | 5.3% | 6.25% | 87.5% | 100% |
| Molestus | 52.6% | 50% | 6.25% | - |
| Hybrids | 42.1% | 43.75% | 6.25% | - |

